# Statistical analysis of a Potts glass model of cortical dynamics

**DOI:** 10.1101/2023.04.05.535702

**Authors:** Kwang Il Ryom, Alessandro Treves

## Abstract

We introduce, in a previously studied Potts model of long-range cortical interactions, a differentiation between a frontal and a posterior subnetwork. “Frontal” units, representing patches of anterior cortex, are endowed with a higher number *S* of attractor states, in keeping with the larger number of local synaptic contacts of neurons there, than in occipital cortex. A thermodynamic analysis and computer simulations confirm that disorder leads to glassy properties and slow dynamics but, surprisingly, the frontal network, which would be slower if isolated, becomes faster than the posterior network when interacting with it. From an abstract, drastically simplified model we take some steps towards approaching a neurally plausible one, and find that the speed inversion effect is basically preserved.

## 1 Local attractors in cortical dynamics

Global oscillations in cortical state, as well as EEG and MEG response patterns, have been approached with linear decomposition analyses, such as spherical harmonics [1, 2, 3]. Yet, it is clear that in order to understand information processing and cognition one has to go beyond such macroscopic descriptions, and consider the local attractor dynamics which has been widely hypothesized to serve as the ubiquitous mechanism for expressing memory functionality [4, 5]. The compartment model by Valentino Braitenberg [6] proposes a separation between local and long range cortical interactions, both of which are envisaged to contribute to associative memory. The Potts associative network [7] can then be construed as an effective model of long-range interactions, that subsumes local ones into the *ad hoc* dynamics of Potts units, with *S* states each, which represent patches of cortex. We have analyzed the *latching* dynamics induced by adaptation and inhibition [8] and discussed how it could play out in free recall paradigms [9].

These studies, however, reduce the cortex to a *homogeneous* network of Potts units, each of which is characterized by the same number of states *S*, positive feedback *w*, time constants *τ*’s for excitation, inhibition and adaptation. This is in contrast with prominent features of cortical organization, which for example point at much higher numbers of local synaptic contacts among pyramidal cells in temporal and frontal, compared to occipital cortex [10], suggestive of a capacity for more and/or stronger local attractor states in the former, or conversely at more linear and prompt responses to afferent inputs in posterior visual cortices [11, 12], suggestive of reduced positive feedback relative to more anterior areas. Other features show gradients that roughly align with these, and all together have been proposed by Changeux and colleagues [13] to define, in particular in the human brain, a *natural cortical axis*. If one attempts to incorporate these features into a *non-homogeneous* Potts network, what are the implications for cortical dynamics? The indications that the dynamics in frontal cortex may be more affected by local attractors need not necessarily imply, it should be noted, an enhanced tendency for individual neurons to be routinely “stuck” in steady states, in which they keep firing at steady rates for a few hundred *msec*. This would be in apparent contrast with extensive evidence for more dynamical forms of coding in frontal cortex, e.g., for changing task contingencies rather than stable visual features [14]; or, moving up to entire populations of neurons and to the human brain, for the encoding of verbs rather than nouns [15, 16] (but see [17]) or of syntax rather than the lexicon [18].

Local attractor states may be composed, only transiently or more persistently, into global attractor states. Studying the dynamics of reactivating such global attractors requires assumptions about the nature and the statistics of the compositionality, and we have analyzed two distinct models in this respect, both for a homogeneous Potts network [19, 20]. Here, however, we want to focus on the dynamics unfolding away from previously acquired global attractors, for example as new attractors are being established, or *learned*. In a learning regime, we expect the lack of *a priori* relations between what has been already acquired and the new compositional representation to be established to turn the cortex, from the point of view of the latter, into basically a disordered system. Do long-range cortical interactions then result into “glassy” dynamics, with critical slowing down and persistent traces of initial conditions? If so, how does the glassy character express itself over the short time scales relevant to cognition? Is it affected by gross inhomogeneities, like the posterior-anterior gradients in cortical parameters mentioned above?

We consider here the most basic and mathematically well defined aspects of these issues, by analyzing a *hybrid* model that integrates in the Potts formulation a crude binary version of the gradient along the “natural” axis (Fig.1); and leaving for later reports more realistic models of cortical dynamics and applications to other domains. As we shall see, even the analysis of what seems like a simple extension of a standard model for an infinite range spin glass reveals some surprising properties.

**Figure 1:**
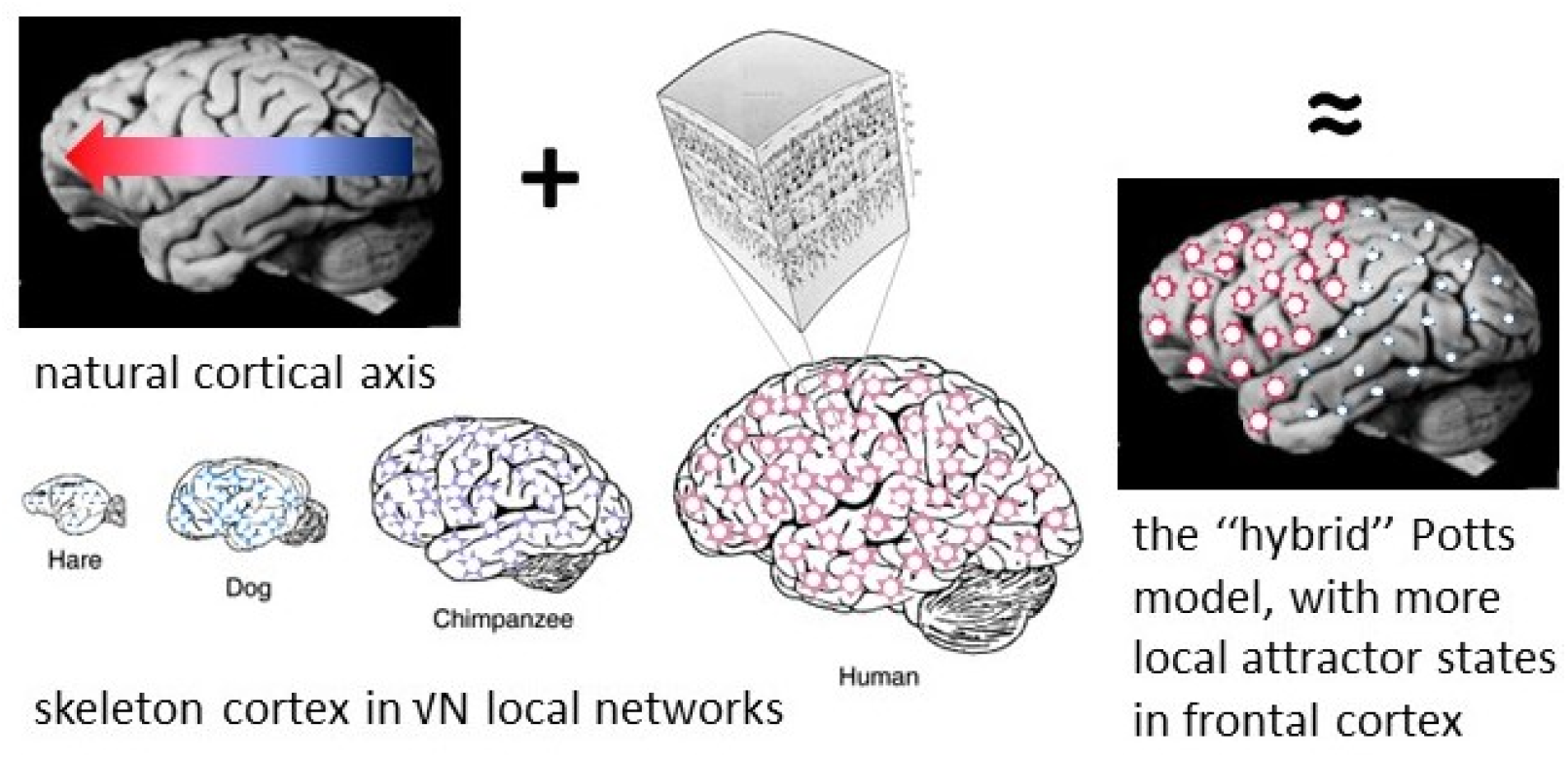
The hybrid Potts model combines the representation of local attractor dynamics in terms of units with *S* active states, inspired by Braitenberg’s idea of an approximate 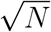 scaling [6], with a differentiation between frontal and posterior cortices, along the natural axis posited by Changeux and others [13] and expressed by a larger *S* value for frontal units. Note the assumption that the critical quantity that varies along the axis is *S*, the simplification of replacing a gradient with just two *S* values, and the ill-fitting temporal cortex areas, in which pyramidal cells have abundant recurrent collaterals [10] but are otherwise included among posterior regions.

## 2 Mean-field analysis of the long-time behaviour

As discussed in previous reports [21, 22], the analysis of the attractor states of associative Potts networks, in which each unit represents a patch of cortex, relies on the same assumption of symmetric interactions, proposed for the standard model [23] in which each unit represents a single neuron.

If we consider, as we do here, a local cortical network to behave effectively as a *discrete* Potts unit, *σ_i_* ∈ {0, 1,…, *S*}, which can take one of *S* active states (labelled by *k* = 1, 2,…, *S*) as well as stay in the quiescent state (labelled by 0), it is convenient to introduce the Potts spin operator,

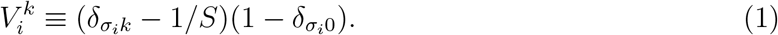

### 2.1 The random homogeneous Potts model, with a zero state

First, we consider a network of Potts units all endowed with the same number of states *S*, that interact through random tensor connections. The Hamiltonian of the system reads

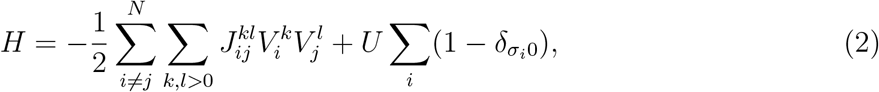

where *N* is the number of Potts units and the 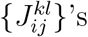 are sampled from Gaussian distributions with mean *J*_0_/*N* and variance *λ*^4^*J*^2^/*N*. We have introduced the normalisation factor *λ*,

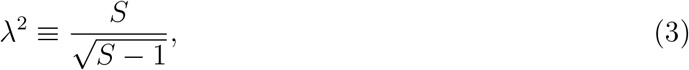

which makes the critical temperature for the transition to a glassy phase independent on *S*, in units of *J* (see below). The interactions satisfy

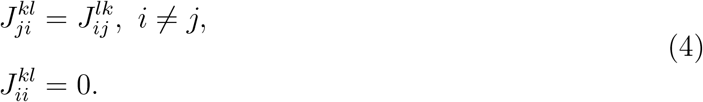

Note that in this model, although *S* is the same across all units, the states of one unit do not correspond to those of another unit, as they would if they represented, e.g., directions in physical space. This is in contrast to the Potts model considered by [24], in which such correspondence holds, and the interactions, albeit still random, are in the form 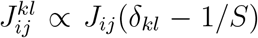, with a single random variable *J_ij_* per unit pair (and, in addition, there is no quiescent state). In that model, the symmetry among Potts states is global, whereas in our model it is local, as it must be in order to represent distinct codes by different patches of cortex.

Despite the larger number of random variables the thermodynamic analysis proceeds along similar lines to that in [24] and it is in some respects simpler. Using the replica method [25], the free energy of the system is written as

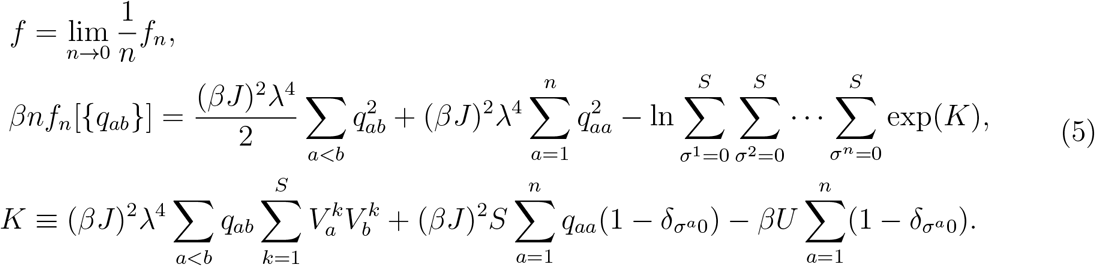

where *q_ab_* is the Edward-Anderson order parameter [25], *β* = 1/*T* is the inverse temperature and replica indices *a* and *b* run from 1 to *n*. The saddle-point equations of Eqs. (5) are

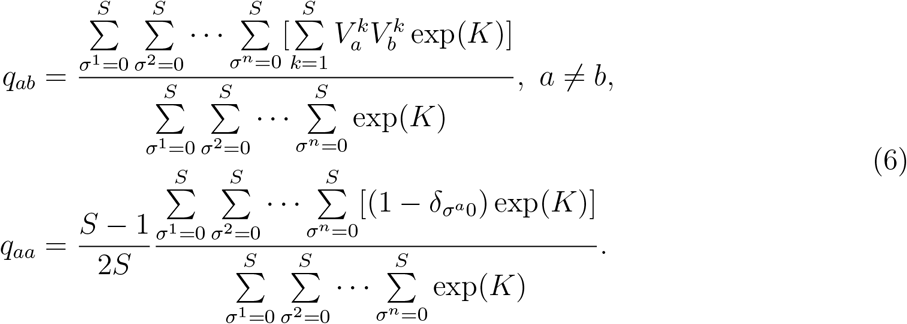

The physical meaning of *q_ab_*(*a* ≠ *b*) is the same as in the Sherrington-Kirkpatrick (SK) model [26] (see also [27]), while 2*q_aa_S*/(*S* – 1) is the fraction of active units in replica *a* of the Potts network. Note that the free energy in Eq. (5) does not depend on *J*_0_, the mean of the normal distribution from which the 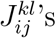 are sampled. This is in contrast with the Elderfield-Sherrington (ES) model [24], where low enough values of *J*_0_ should be chosen to avoid ferromagnetic ordering at low temperatures ([24, 28]). Since the symmetry in our model is local — a sort of *gauge* invariance – there is no meaning to ferromagnetic alignment.

#### Properties near the critical temperature

The paramagnetic solution (*q_ab_* = 0, *a* ≠ *b*) is the ground state of the system at high enough temperatures. Lowering the temperature, a phase transition from the paramagnetic to the spin glass phase occurs at *T* = *T_c_*. To determine *T_c_*, one can (Landau-) expand the free energy close to it, assuming *q_ab_*(*a* < *b*) to be small, to find

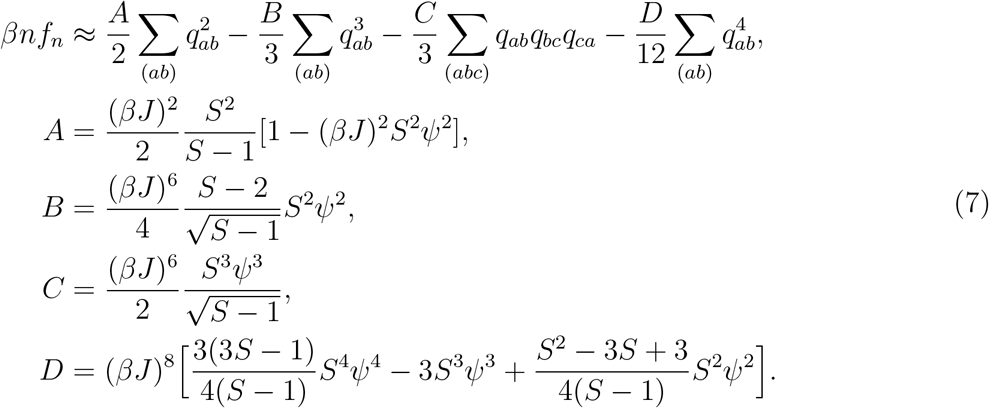

Here (*abc*) means that replica indices *a, b, c* are all distinct in the summation. Following [29], we have retained only the quartic term that is relevant for replica-symmetry breaking (RSB) in Eqs. (7). We have also assumed that the order parameter *q_aa_* does not depend on the replica index *a* near *T_c_* and thus have introduced a symbol *ψ* ≡ 2*q_aa_*/(*S* – 1) to reduce the burden of heavy notation. Then, *ψ* should satisfy

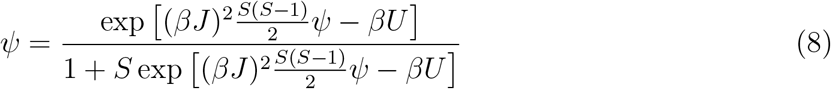

and the quantity *Sψ* gives the fraction of active units in the network.

Under the replica symmetric (RS) assumption, *q_ab_* = *q* (*a* ≠ *b*), the critical temperature is determined by numerically solving

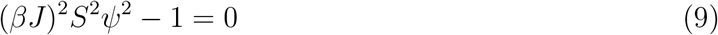

together with Eq. (8), since *ψ* contains *T*. If *U* → –∞, then *Sψ* → 1 and we get a simple formula, *T_c_* = *J*. The phase transition is a continuous one if

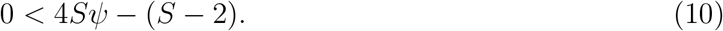

For *S* < 6, there exists a critical value of *U, U_c_*, above (below) which the transition is discontinuous (continuous). For *S* > 6, the transition is discontinuous for all values of *U*. This RS solution is however unstable against RSB in the whole glassy phase. Thus, replica symmetry should be broken.

To probe replica symmetry breaking, following Parisi’s hierarchical scheme [30] we write the free energy

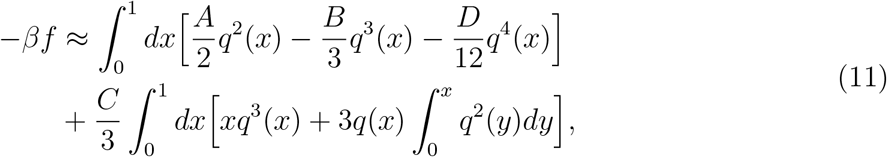

which is to be maximised with respect to Parisi’s function *q*(*x*) [29]. We note that Eq. (11) has the same form as in the ES model, except for the coefficients. Thus, we can envisage that the nature of RSB is similar to that in the ES model (see also [28, 31] for its detailed properties). The non-trivial solution of Eq. (11) is

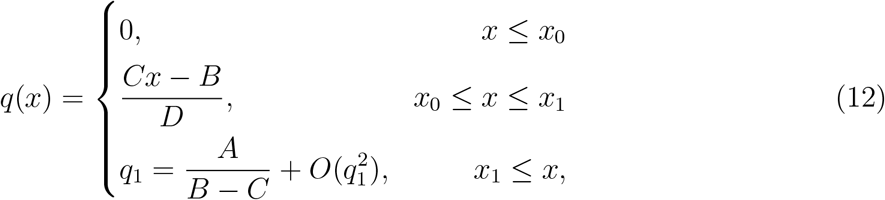

where

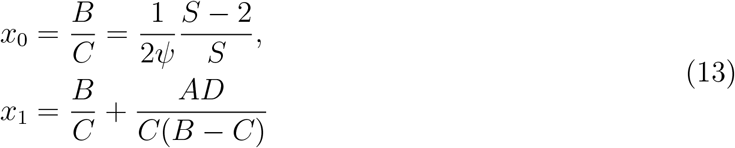

From Eq. (12), we can see how replica symmetry is broken, for a given value of *U*. The scheme in Fig.2 is similar to the one for the ES model. Note that *x*_0_ is always zero for *S* = 2, regardless of *U*, whereas it remains positive for *S* > 2. This means that *P*(*q*) has a Dirac delta at *q* = 0 for *S* > 2, whereas there is no Dirac delta at *q* = 0 for *S* = 2, as in the SK model. The phase transition to the glassy phase is continuous if

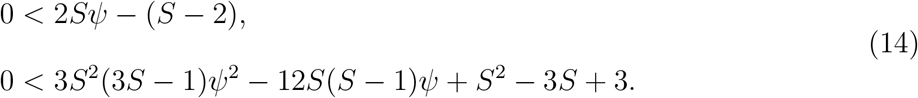

In general, these two conditions are numerically probed together with Eqs. (8) and (9) for a given value of *U*. As a special case, when *U* → –∞, the second condition is guaranteed.

**Figure 2:**
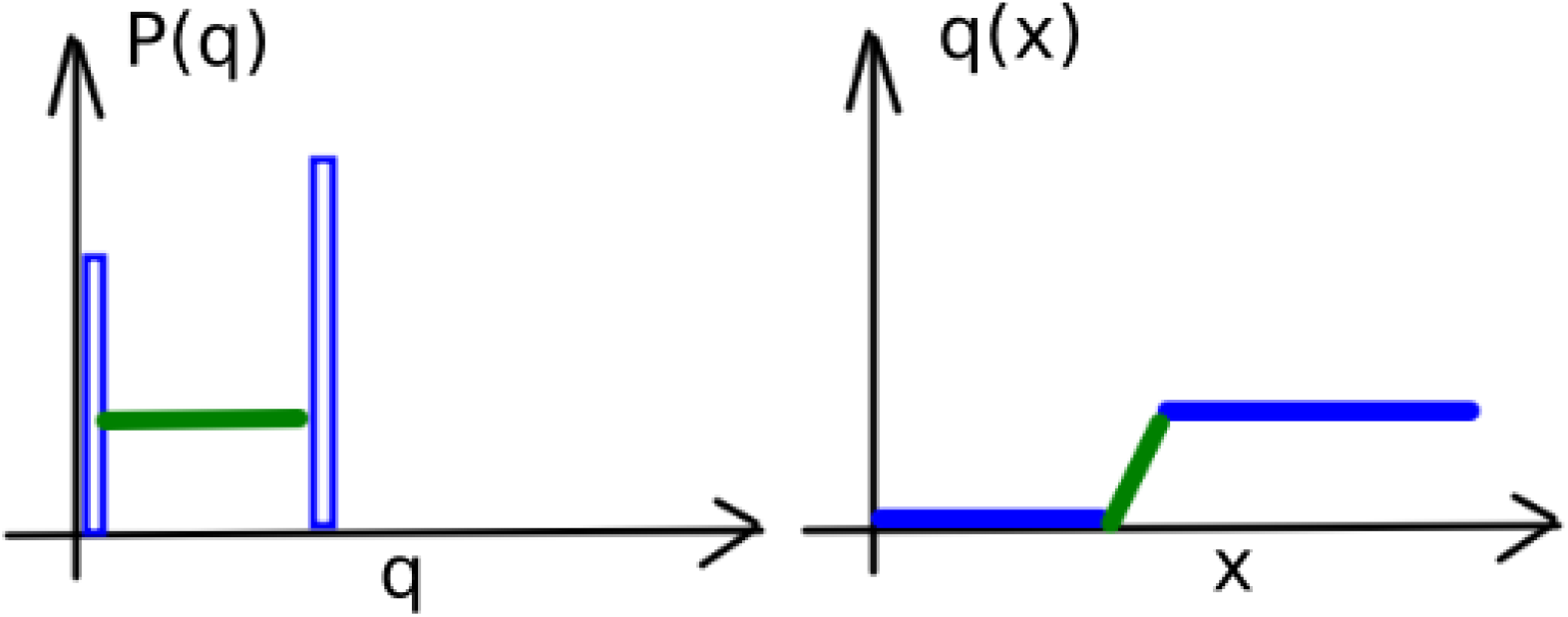
Schematic description of replica symmetry breaking, from Eq. (12). Left: the probability density *P*(*q*), with blue rectangles denoting Dirac delta functions. Right: Parisi’s function. Colour coding is used to facilitate a visual comparison.

However, unlike the RS Eq. (10), the first of RSB Eqs. (14) ceases to hold for *S* > 4. Thus, the transition can be continuous only for *S* ≤ 4. We can compute the range of *U* where Eqs. (14) hold by solving them together with Eq. (8) and Eq. (9). The result is shown in Fig.3(a).

**Figure 3:**
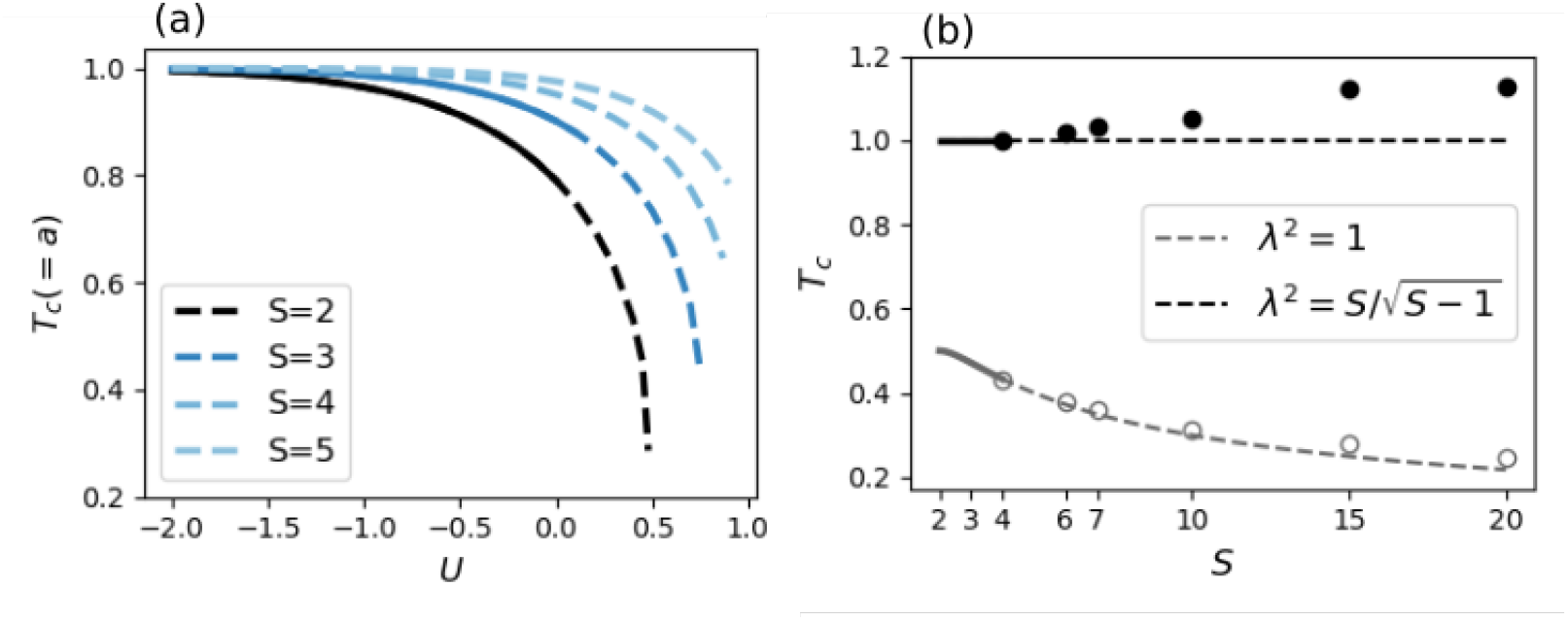
The critical temperature. (*T_c_*) for the onset of the glassy phase of a homogeneous Potts network. **(a)**: *T_c_* as a function of the threshold *U* for a model with a zero state. With the normalisation set as in Eq. (3), the mean activity *a* of the network at *T* = *T_c_* is equal to *T_c_* itself (that is, to *T_c_*/*J*). Dashed curves are predicted by RS theory and solid curves are from RSB theory. All transitions shown here are continuous. **(b)** *T_c_* as a function of *S* for a model without a zero state (*U* → –∞): colour encodes the normalisation used (as indicated in the legend). Solid curves are obtained analytically from the Landau expansion of the free energy (a continuous phase transition) and dashed curves are their mere extensions, to guide the eye. Circles are obtained by numerically maximising the 1-step RSB free energy (a discontinuous transition), Eq. (16). We set *J* = 1.

#### Properties at all temperatures

At temperatures well below *T_c_* and when the transition is discontinuous, one should directly deal with the free energy, Eqs. (5), in the full RSB formalism. Even for the SK model, solving Parisi’s equations requires sophisticated numerical techniques (see [32]). However, Potts spins seem to have a distinguishing property from Ising spins, at least when we compare the ES model with the SK model: while any finite-step RSB solution is unstable in the SK model ([30]), the first-step RSB (1RSB) solution is locally stable in the ES model (*S* > 2) below *T_c_*, down to a certain temperature, where another phase transition occurs [28, 31]. So, one can study discontinuous transitions for *S* > 4, where the Landau expansion does not apply, by using a 1RSB formalism ([33]). Below we use this method to study the discontinuous transition of our random Potts model, Eq. (2).

While the expression of the free energy Eq. (5) within a 1RSB ansatz is given in the Appendix (see Eq. (40)), its numerical solution is computationally hard, especially for large values of *S*. Thus, we restrict ourselves to a special case: the threshold *U* goes to –∞ (the zero state then drops out of the equations). Inspired from the shape of Eq. (12), we seek solutions of the form

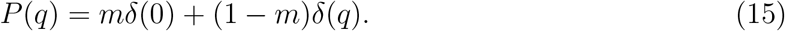

Then, the 1RSB free energy becomes,

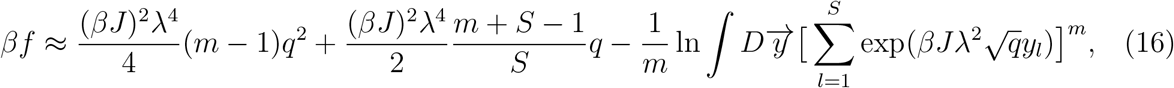

where

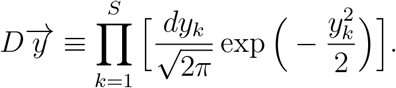

We can numerically maximise Eq. (16) by using the same numerical trick as in [33], up to *S* = 20. Critical temperatures obtained that way are reported in Fig.3(b), while the order parameters are shown in Fig.4.

**Figure 4:**
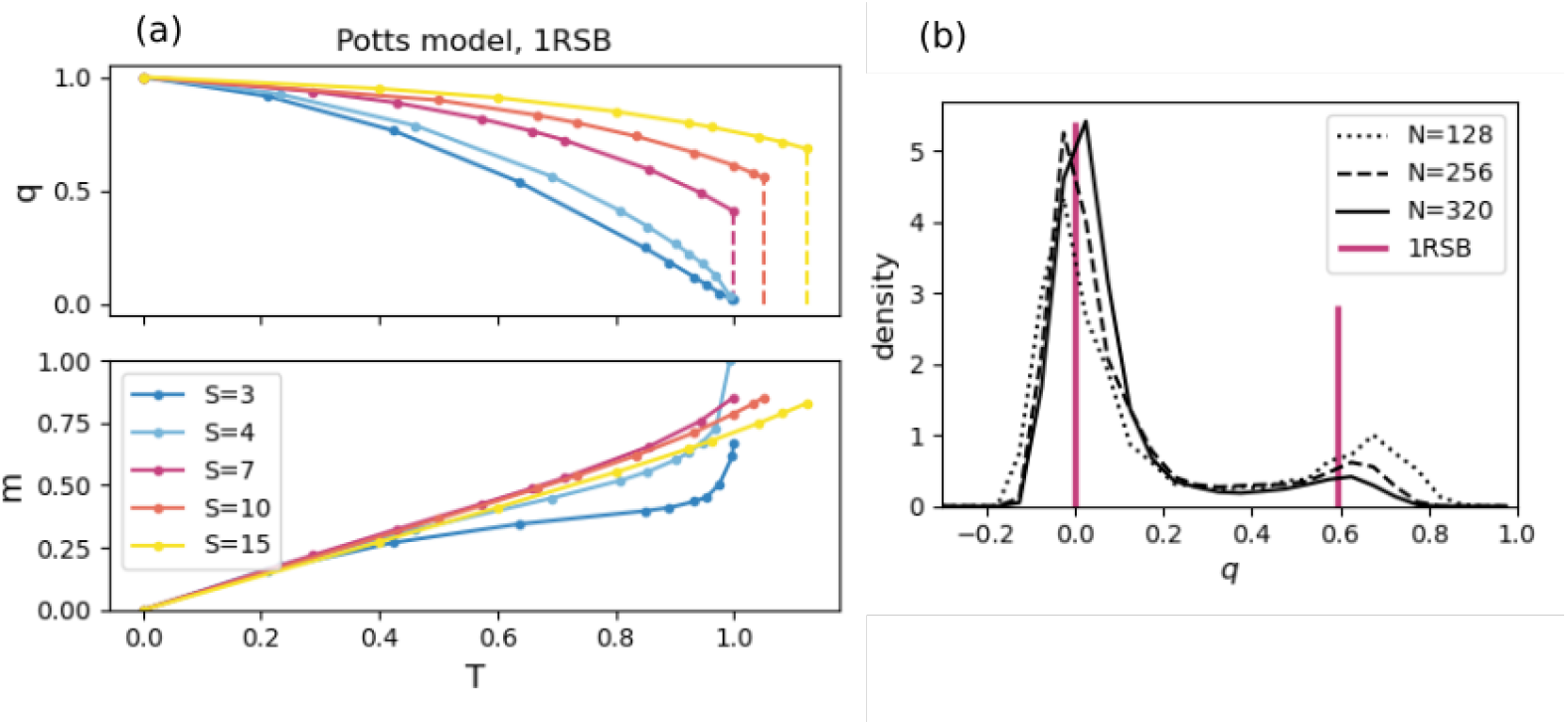
Order parameters of a homogeneous Potts network without a zero state (*U* → –∞), predicted by 1RSB theory. **(a)**: solutions of the 1RSB free energy as a function of *T*. Note the discontinuous jumps in *q* at *T* = *T_c_* for *S* > 4. **(b)**: Probability density, *P*(*q*), obtained from Monte Carlo simulations, for *S* = 7 and 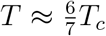. Red vertical lines indicate Dirac delta functions, estimated from (a). The peak at higher *q* seems to be lower with increasing values of *N*, but this is due to the insufficient relaxation time in our simulations. Since the relaxation time grows exponentially with *N* ([34]), we did not attempt to obtain the exact ground states.

### 2.2 The hybrid Potts model without a zero state

We now consider a network of Potts units that have different values for *S*: a unit *i* has its own number *S_i_* of Potts states. For the sake of simplicity, we consider Potts units without the quiet state. We group units according to their number of states: there are *N_l_* units in group *l* (*l* = 1, 2,…, *L*) and they have *S_l_* Potts states each. If the total number of Potts units in the network is *N*,

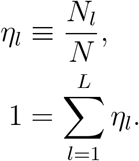

We write

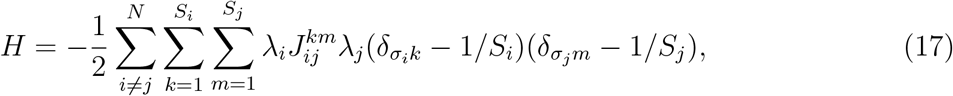

where 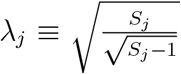 normalizes the interactions with both a pre- and a post-synaptic factor, and the 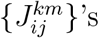 are sampled from a Gaussian distribution of mean *J*_0_/*N* and variance *J*^2^/*N* and satisfy Eqs. (4).

The replica method gives the free energy,

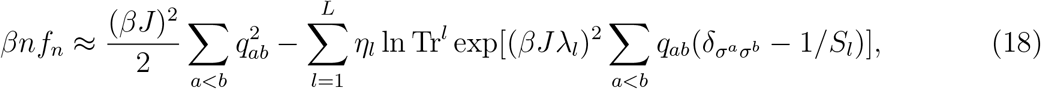

where *q_ab_* is the Edward-Anderson order parameter,

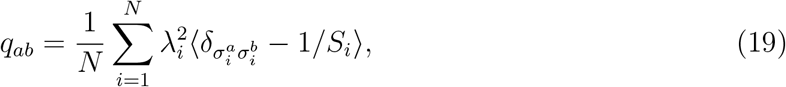

and

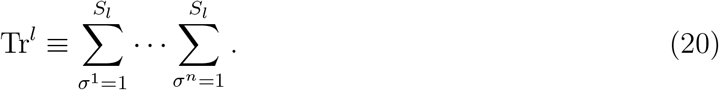

As in the previous subsection, we expand Eq. (18) around *q_ab_* = 0 and apply the Parisi algebra [35] to probe the nature of the equilibrium state. The corresponding free energy functional and the Parisi function that maximises it have the same form as for the homogeneous network (see Eqs. (7), (11), (12)), after a redefinition of the coefficients *A, B, C* and *D*

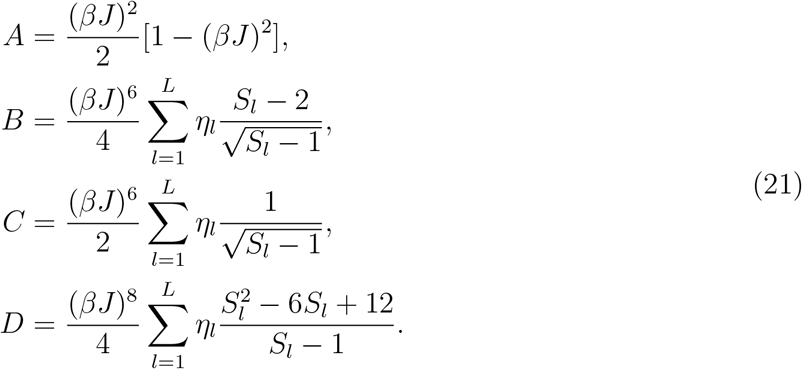

The critical temperature for the onset of the glassy phase is again given by

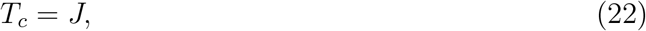

where the phase transition is continuous in terms of *q* whenever

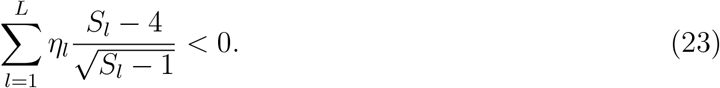

As an example, consider a hybrid network with two types of Potts units: half with *S*_1_ and half with *S*_2_ states. The phase transition is continuous if

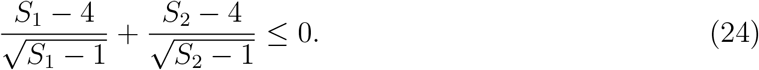

Several cases are interesting (we set 1 < *S*_1_ ≤ *S*_2_):

- *S*_1_ = 2. The transition is continuous for *S*_2_ ≤ 10 and discontinuous otherwise.
- *S*_1_ = 3. The transition is continuous for *S*_2_ ≲ 5.5 and discontinuous otherwise.
- *S*_1_ ≥ 4. The transition is always discontinuous (except for *S*_1_ = *S*_2_ = 4, but then the network is again homogeneous, as in the previous section).

### 2.3 The glassy phase of a Potts associative memory

We consider now an attractor neural network comprised of Potts units. The Hamiltonian is the same as in Eq. (2), with the connection 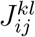 now given by the Hebbian-learning rule,

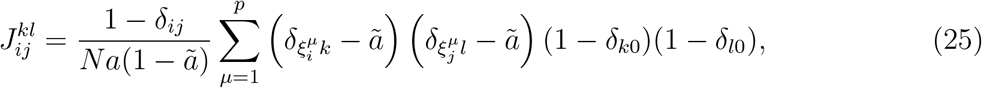

where *ã* = *a*/*S* and 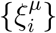 are *p* randomly correlated memory patterns. The free energy is obtained by the replica trick (see [36] for the Hopfield model and [21, 22] for the Potts model):

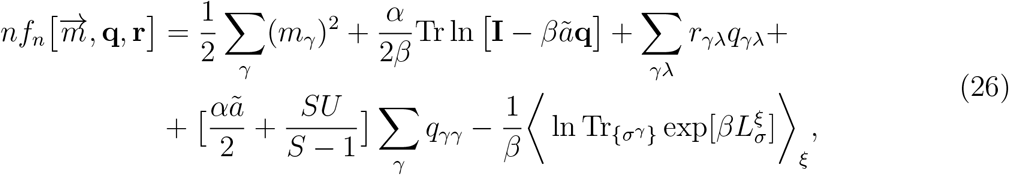

where

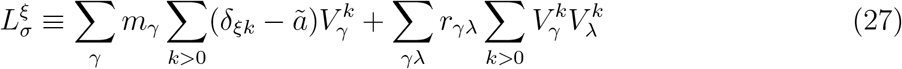

and *α* ≡ *p*/*N*. The saddle-point equations read

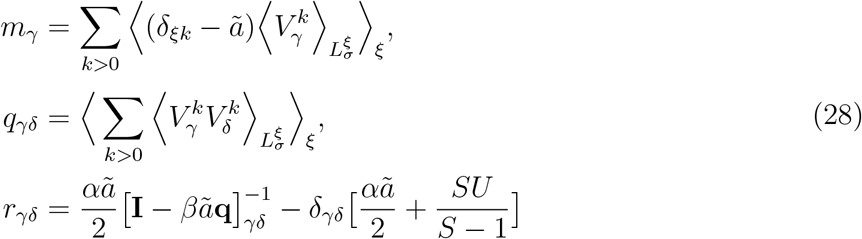

where

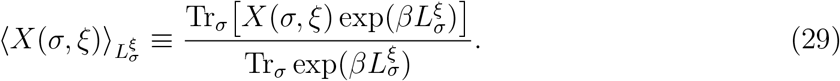

One can solve Eqs. (28) by using either RS or RSB assumptions to compute, *inter alia*, the storage capacity of network. We refer to Refs. [21, 22] for a discussion of the storage capacity (see also [7, 37] for related but different models). Here, we are interested in the phases prevailing at higher temperature: the paramagnetic and the glassy phase.

At high enough values of *T*, in fact, retrieval solutions do not exist. So, we set *m_γ_* = 0 and the terms including *ξ* and *m_γ_* drop out of the equations. We can easily see that *q_ab_* and *r_ab_* converge to zero in the high temperature limit, if *a* ≠ *b*. We expand the free energy with respect to these two variables around zero

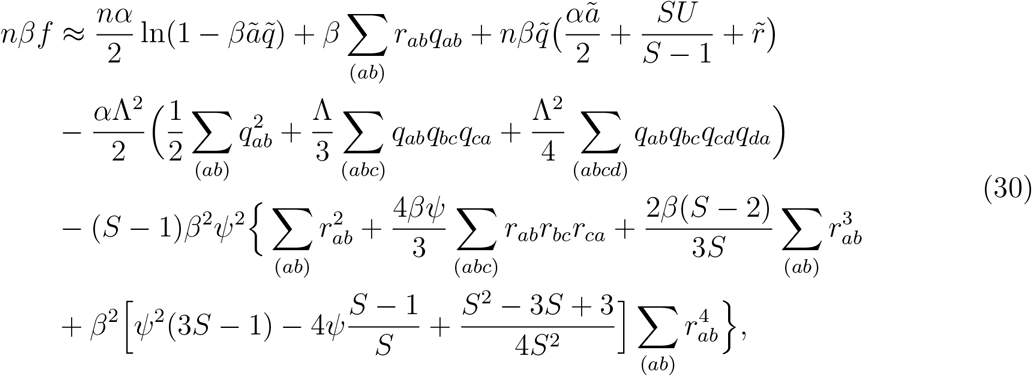

where

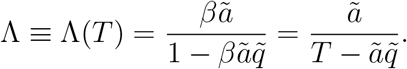

For the sake of simplicity, let us consider a RS ansatz. Then, the free energy reads, up the third order in *q* and *r*,

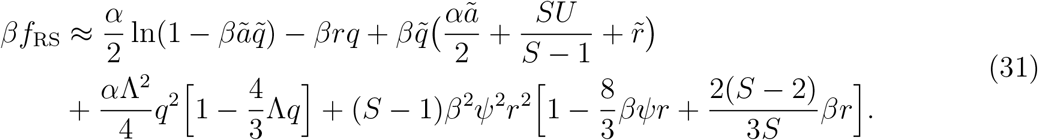

This free energy is maximised with respect to *r* and *q*, while 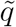 and 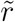 satisfy

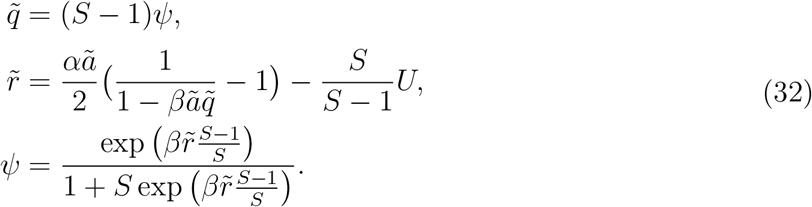

In addition to the trivial (paramagnetic) solution of *q* = 0, we have

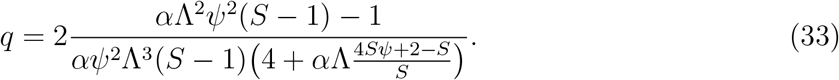

A phase transition from the paramagnetic to the glassy phase occurs at

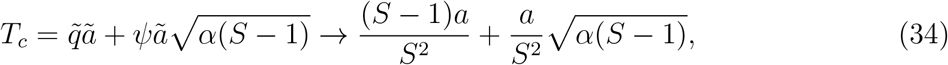

where the last expression is for the limit of *U* → –∞. It is a continuous transition if *S* ≤ 6.

For *S* > 6, the transition is continuous if *α* < *α*_0_ and discontinuous otherwise, with

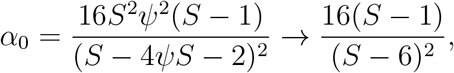

where the last expression is again for *U* → –∞.

As in the the random Potts model considered above, there is a value of *U* above which the phase transition cannot be treated by the Landau expansion (either the phase transition does not occur or it cannot be described by the Landau expansion). This result is shown in Appendix.

## 3 Dynamics

Although dynamics can be studied within mean-field theory to a certain extent ([38]), here we stick to Monte Caro (MC) simulations. Throughout this work, we use the heat bath algorithm to simulate the dynamics of Potts networks. Specifically, the local field of each Potts unit is computed as

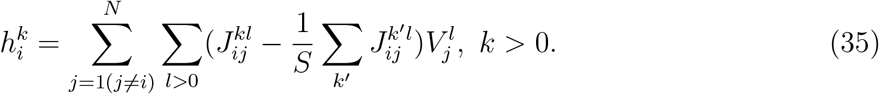

At each MC step, one Potts unit is randomly chosen to be updated based on the following equations

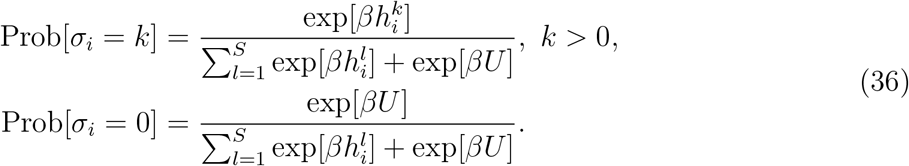

For models without a zero state, the second of Eqs. (36) is not used.

For most of the simulations presented here, we run two systems ([34]) with the same quenched disorder (i.e., the set of interactions between Potts units) and measure the overlap between the two configuration at time *t*:

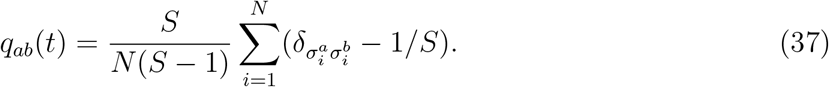

### 3.1 Dynamics close to steady states

Fig.5 shows sample trajectories at temperatures *T* ≪ *T_c_*, to illustrate their glassy nature: after an initial transient the system is trapped in metastable states for a while before finding a way out, along which it can further lower its energy. The opportunities to escape a metastable state however become rarer and rarer, and the time spent near it longer and longer, a process called thermalization.

**Figure 5:**
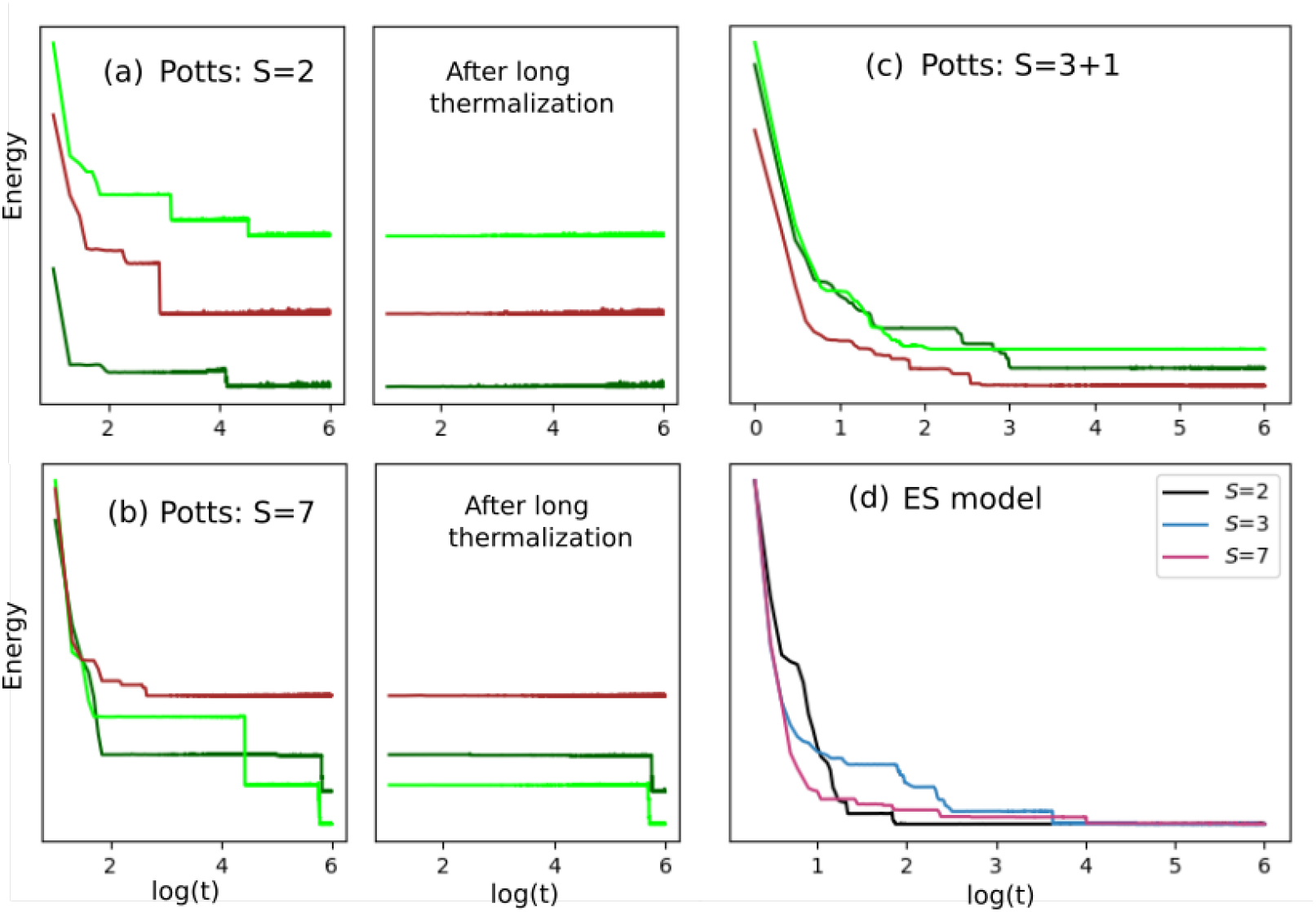
Energy as a function of MC sweeps per unit for sample MC trajectories. Note the log scale of the abscissa. **(a)**and **(b)**: three example trajectories are shown for a homogeneous Potts network without a zero state. In the right panels, *t* restarts after *t*_0_ ≃ 10^5^, to focus on long-time glassy dynamics. **(c)**: example trajectories of a homogeneous Potts network with a zero state. **(d)**: example trajectories of the ES model ([24]). The three curves are rescaled and shifted for better visibility (only in panel (d)). Note that the ES model reduces to the SK model if *S* = 2. The number of units is *N* = 256 for all three panels, and each data point is averaged over 10 MC sweeps, except for the first 100 points.

To measure how fast the dynamics unfolds on the glassy free energy landscape, we first “thermalize” a configuration by letting it evolve for *t*_0_ = 10^3^ time steps, and then start from it two simulations with identical interactions, until at *τ* their overlap reaches half its initial value.

Since the times τ are quite scattered depending on the realization of the interaction – their logarithms are approximately normally distributed – we consider their cumulative distribution, for a given network, and in particular the thermal *half-life* scale *ζ_g_*(*T*), defined as the median *μ*_1/2_[log(*τ*)] when the cumulative distribution, at a temperature *T*, reaches the value 0.5.

With this procedure, we find that a homogeneous random Potts network “moves” faster the smaller are its Potts units, i.e., the lower their *S*. This is shown in Fig.6a, which indicates that *ζ_g_*(*T*) ≡ *μ*_1/2_[log(*τ*)] increases approximately with log(*S*), with the parameters we use.

**Figure 6:**
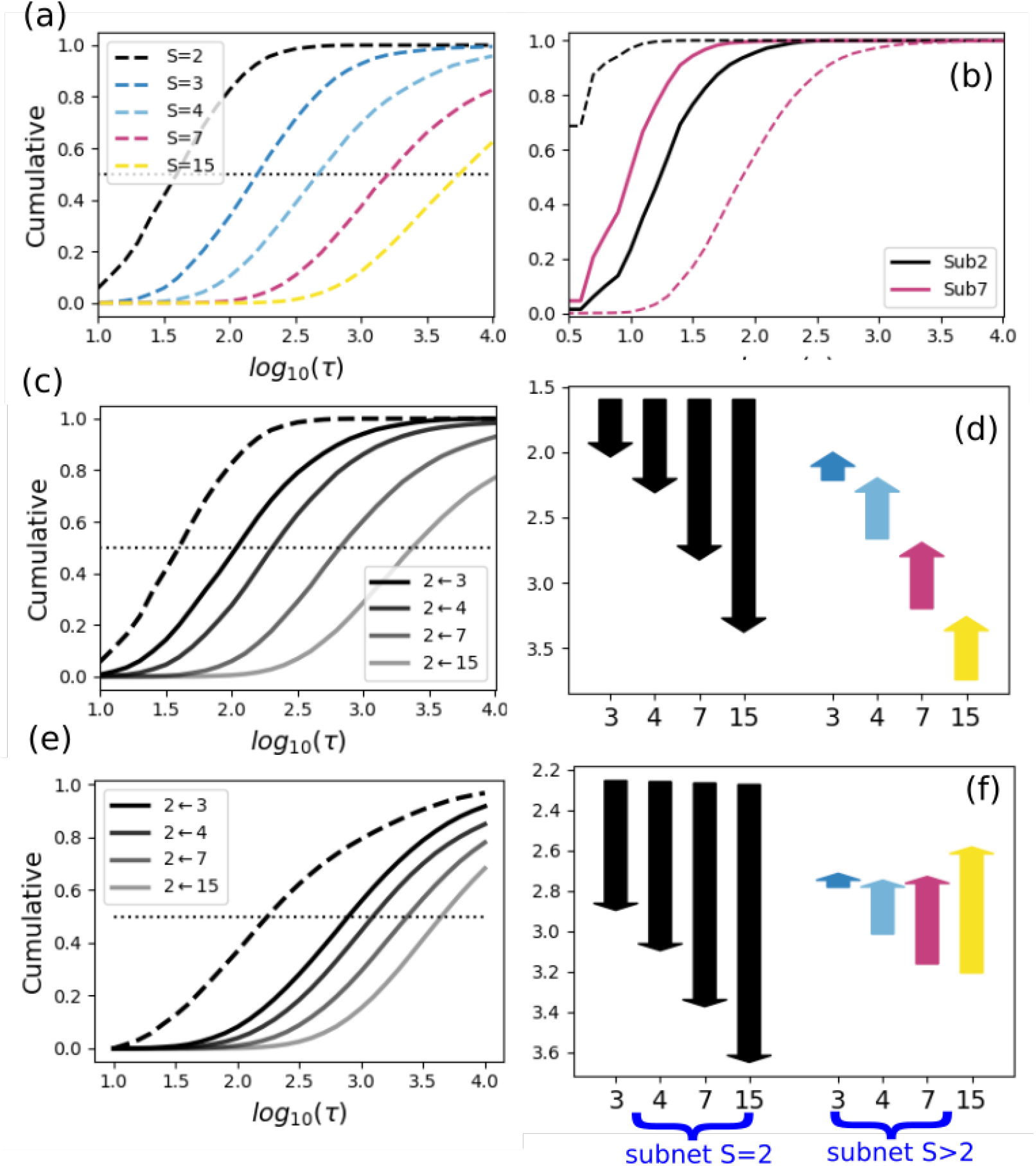
Speed-up and slow-down in a hybrid Potts model. **(a)**: Cumulative distribution of *τ*, computed for homogeneous networks of *N* = 256 units, as a function of *S*. **(b)**: the inversion of speed due to hybridization between spins (*S*_1_ = 2) and large units with *S*_2_ = 7. **(c)**:a sub-network of *S*_1_ = 2 that interacts with another sub-network with *S*_2_ > 2 is more slowed down the higher is *S*_2_. Note that the case with *S*_2_ = 2 is the homogeneous network of panel (a). **(d)**: The speed-up and slow-down of sub-networks are shown by the arrows, which head up for units that accelerate. The color of bars stands for S as in panel (a), while the height measures the difference Δ*ζ_g_* in the median of the cumulative distribution of *τ*, between hybrid and homogeneous networks. **(e)** and **(f)**: same as (c) and (d), but without the normalization constants *λ_i_* in Eq. (17) and *T* = 0.2, *t*_0_ = 5 × 10^3^.

If we measure *τ* (and *ζ_g_*) separately for the units with a given *S* in a hybrid network, we find that the small units get slower and the large units get faster, due to the hybridization. Surprisingly, however, the effect is not simply an interpolation or averaging of the temporal scale between the two subnetworks, that would come to share a common speed, because in many cases the large units get markedly faster than the small units. This is shown in Fig.6b for *T* = 0.8 and large units with *S*_2_ = 7, that interact in a hybrid network with small spins, *S*_1_ = 2. Fig.6c shows that the slowing down of these spins scales roughly with the log of *S*_2_, the number of states of the units that “bog them down”. Simultaneously, the large units “speed up” after the hybridization, Fig.6d and, particularly when the interactions are not renormalized as in Eq. (17), can get to be faster, on average, than the small spins (Fig.6f).

The speed inversion phenomenon indicates that the same free-energy landscape is “perceived” as rougher, near the metastable states, by Potts units with fewer degrees of freedom. Does the same effect occur away from the metastable states, e.g. in the initially rapid dynamics to the left of the panels in Fig.5, or when *asymmetric* connection weaken the very stability of such disordered states?

### 3.2 Factors that accelerate the dynamics

In the Hopfield model, imposing symmetry in the interactions, which established the connection with Hamiltonian systems, thus enabling the analytical approach [23], entailed gross disregard for Dale’s law – stating that excitatory and inhibitory neurotransmitters are released by distinct types of cortical neurons – and also of plausible statistical models of connectivity among excitatory neurons alone. Interestingly, it was argued early on that spin-glass-like metastability would still characterize networks with asymmetrically “diluted” connectivity, whereas it was suggested that more profound changes due to asymmetry might be observed in the dynamics [39]. In the Potts network, inspired by Braitenberg’s model [6], Dale’s law is not relevant as long-range connectivity (the component modelled by the tensor interactions among Potts units) is only excitatory; and there is no urgency to consider diluted connectivity either, as the tensor connections themselves are considered to recapitulate thousands of individual synaptic connections [22]. Still, it makes sense to consider the effect of asymmetric non-zero values in the random interactions, by writing them in the form

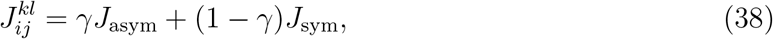

where 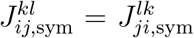 and 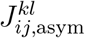 is unrelated to 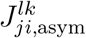 thus the former are *symmetric* and the latter *a-symmetric* random components, drawn from the usual distribution with zero mean and variance *J*.

Fig.7b,c shows that introducing asymmetry does have a major effect in speeding up the dynamics, across the board, while maintaining the slowing down of small units and speeding up of large units due to hybridization. With *γ* = 0.3, the root-mean-square symmetric component of the weights is still more than twice the asymmetric component, and yet dynamics are extremely fast.

**Figure 7:**
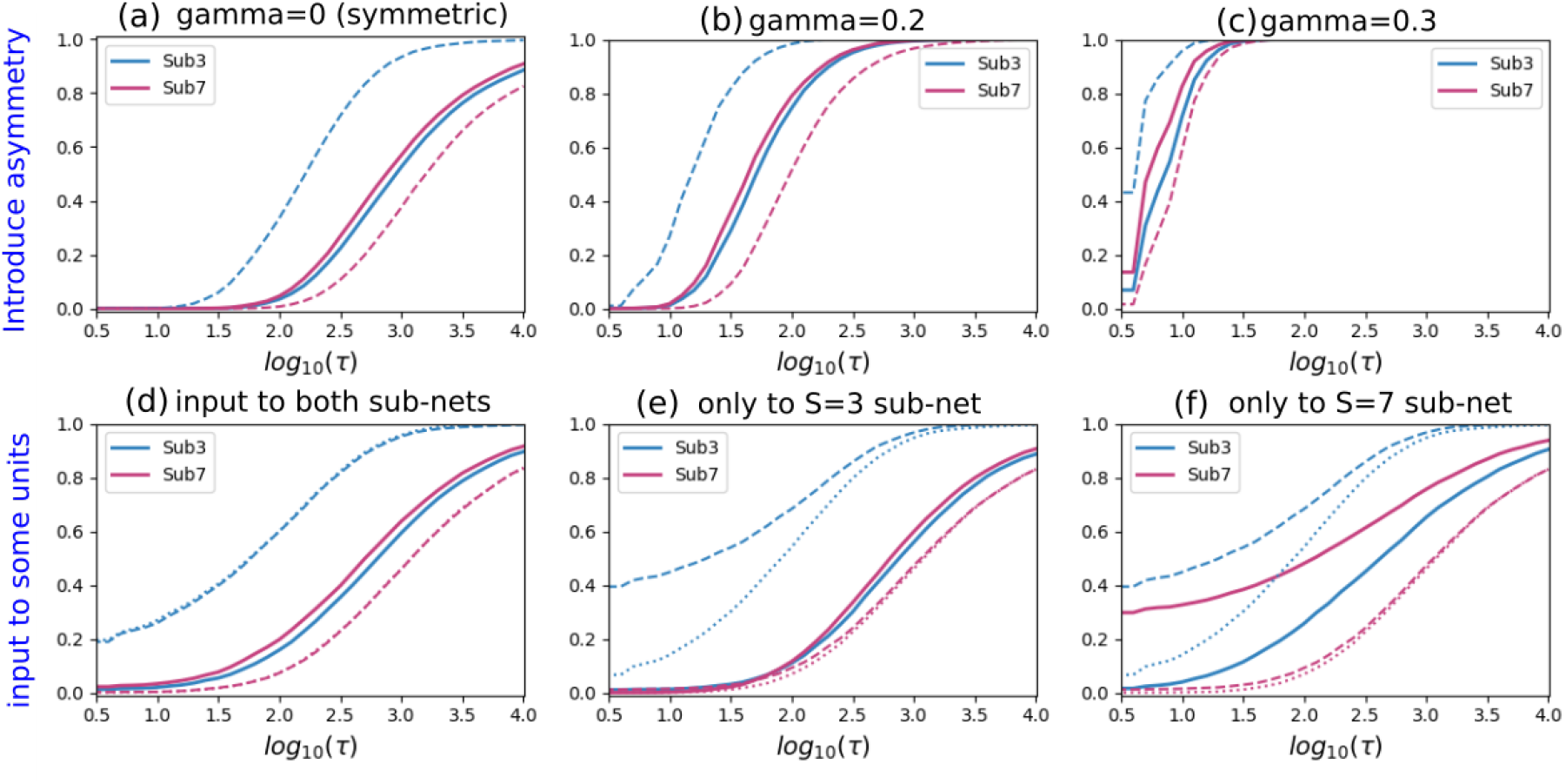
Speeding up the dynamics with asymmetric connections and external inputs. **(a)**: The speed-up and slow-down of sub-networks (relative to their homogeneous counterparts) are shown without asymmetry or perturbation, to serve as the “control” case. **(b)**: The effect of asymmetry, where *γ* = 0.2, is to speed up the dynamics across both subnetworks, homogeneous or hybrid. **(c)**: With more asymmetry, *γ* = 0.3, the same general speed-up is seen as in (b), but more extreme. **(d)**: *N*/4 units are perturbed or reset, after thermalization, mimicking an external input; they are selected uniformly across the whole network. **(e)**: Those perturbed by the input are all in the smaller unit sub-network. **(f)**: They are all in the larger unit sub-network. In both (e) and (f) the dotted curves refer to unperturbed halves of homogeneous networks, and the dashed ones to the halves including the units receiving the input.

To probe the dynamics away from the vicinity of the metastable states, without touching the symmetry of the interactions, we use a variant of the simulation paradigm above, that mimics the arrival of an external input to the Potts network. That is, after a configuration has been thermalized as in previous simulations, a fraction of the units are randomly reset in a new state (different from the thermalized one), and then two independent trajectories evolve with the heat bath procedure from this common starting configuration, until the time *τ* when their overlap has been halved. Fig.7 shows that the basic inversion effect, and in particular the selective slowing down of the “small” units, persists over wider regions of activity space. With respect to the standard thermalization paradigm in Fig.7a, panel d shows that resetting a quarter of the units does indeed accelerate the dynamics of the *S* = 3 network, when it is homogeneous; whereas when it is hybridized with *S* = 7 units, these latter get faster, and slightly faster than the *S* = 3 ones.

In Fig.7d, the external stimulus or perturbation is applied to a quarter of the units distributed in both sub-networks; when they are concentrated among the small *S* = 3 units (panel e, solid curves), the already minimal acceleration effect is reduced even further. When they are concentrated among the large *S* = 7 units, instead, their sub-network activity is markedly accelerated (panel f, solid curves), but only if it is part of a hybrid network with *S* = 3 units, with only minimal acceleration if they are part of a homogeneous network.

The results of the simulated external input procedure are therefore rather counter-intuitive: if affecting one fourth of the Potts units, the input effectively distances the network from its slow-evolving glassy state in two situations: when it is applied to a homogeneous network of small, but not large units, *or*, in a hybrid network, only when it is applied to the large units, but then it accelerates essentially their dynamics alone. These complex effects are observed still within the domain of networks with symmetric interactions, and they beg the question of what happens when an external input is combined with relaxing the symmetry constraint in a cortically plausible manner.

### 3.3 Approaching a cortical scenario

An interesting model of how cortical dynamics might influence cortical connectivity might be expressed by setting *γ* = 0 only for the interactions among the small units. This leads to a remarkable inversion effect, illustrated in Fig.8a. One can see a self-consistent pattern potentially at play: the hybridization makes the large-S subnetwork fast, which upon synaptic plasticity would tend to result in more asymmetric tensorial couplings connecting those units.

**Figure 8:**
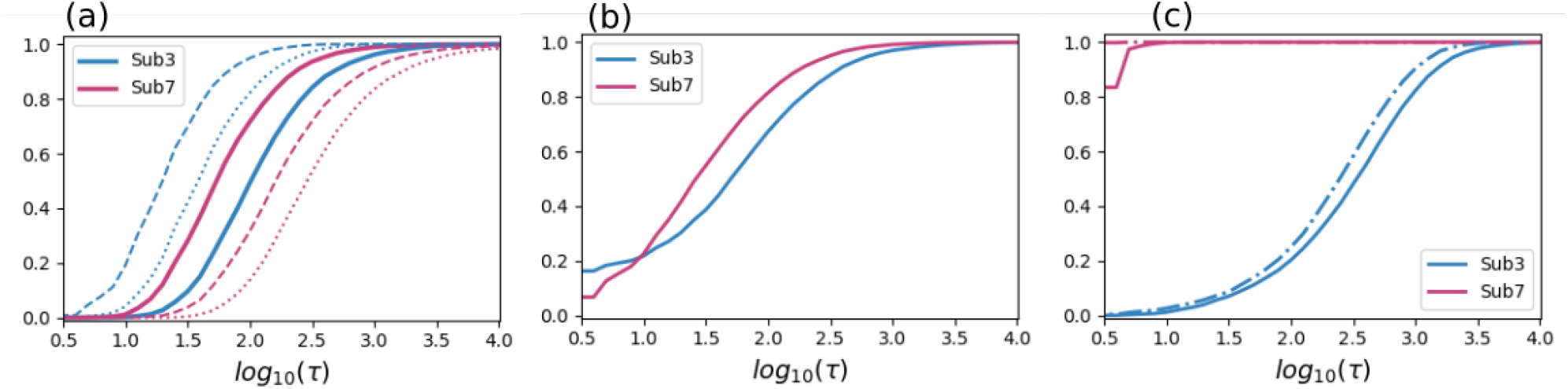
The speed inversion effect likely applies to the cortex. **(a)**Distribution of divergence times when the asymmetric component is zero only within the *S* = 3 sub-network and *γ* = 0.2 otherwise. For homogeneous networks, dotted curves are for the sub-networks that have zero asymmetric component. **(b)** Same as in (a), but half the *S* = 3 sub-network units are perturbed after thermalization. **(c)**. Potts glass model with a quiet state and with regulated mean-activity. After thermalization, a persistent external input is applied to the *S* = 3 sub-network, by flipping to a different active state a proportion *aη* of its active units, inactivating a proportion (1 – *a*)*η*, and activating (in a random active state) the exact same number as those that get inactivated (which is close to *Na*(1 – *a*)*η*, but varies somewhat in the course of each thermalization). The newly activated units are clamped. Broken curves show results when reintroducing asymmetry, *γ* = 0.2, in the connections involving the *S* = 7 units.

To combine a putative external sensory input and the same type of asymmetry of Fig.8a, in a cortically plausible scenario, we show in Fig.8b what happens when resetting a fraction *η* of the small-*S* units (thus simulating an input to posterior cortex) after thermalization. The result is a moderate general speed-up, for both sub-networks, and very fast dynamics in about 30% of the runs, for the posterior network. It appears that in those runs the input has brought the small-*S* units close to the boundary between deep basins of attraction, so that fast noise leads to the immediate divergence of trajectories with the same starting point. For most of the other runs, instead, presumably well inside each large basin, the posterior network remains slower than the frontal one.

Finally, in Fig.8c we take a major step towards cortical plausibility, by re-introducing the quiet state until now considered only in the thermodynamic analysis. The quiet state implies sparse activity (only a fraction *a* of the Potts units in one of their active states) and this overall level of sparsity must be conceived as being regulated by inhibition (in the analysis, this amounts to considering the activity level rather than the threshold *U* as a parameter, whereas for the implementation in the simulations see the Appendix). We first consider in this case purely symmetric random connections, and an input applied to some of the posterior units. To maintain the sparsity level, the input is applied after thermalization both to units already in an active state (which are then flipped to a different state) and to units in their quiescent state – in this case the input is *clamped* to keep them in the new state, simulating the strong effect of thalamic inputs impinging on an inactive local network. Again, we refer to the Appendix for a full description of the procedure. The result, in Fig.8c is a strong differentiation between slow dynamics in the posterior network and an immediate divergence of nearly all trajectories in the frontal one. While this outcome stems to a large extent from clamping a few critical units only in the posterior network, it suggests that the main speed inversion phenomenon is not necessarily reversed back again when moving towards actual cortical dynamics. Reintroducing the asymmetry in the connections involving the *S* = 7 units only makes their network diverge immediately in *all* trajectories (the broken curves in Fig.8c).

### 3.4 Short-term dynamics for the associative memory model

In this last Results subsection we consider the associative memory model, in which the interactions are not random but rather tend to align the network along one of a number *p* of pre-acquired memory states. Here there is no hypothesis about the overarching structure of memory representations in the cortex (we have reported elsewhere on the problems in applying to the cortex the simplest autoassociative retrieval scenario [20]) but rather we aim to to assess the effects on glassy dynamics of the presence of the large attractors associated with the memories. The logic is that we are probing the establishment of new representations, driven by either external inputs or internal dynamics, and if the network gets stuck into a previously acquired memory, no new configuration can be learned.

First, Fig.9a shows that hybridization, i.e., the differentiation between large- and small-S units, in this case speeds up both subnetworks. In a homogeneous network, the *S* = 7 units are extremely slow, as nearly all trajectories are trapped in one of the large basins of attraction of the memories encoded in the connections, reflecting the very extensive storage capacity of the Potts network, quadratic in *S* [7, 21]. Also the trajectories of the *S* = 3 homogeneous network are slower than in the random network, which does not have the memory attractors, but faster than the *S* = 7 ones. The effect of hybridization is then much stronger on the *S* = 7 units.

**Figure 9:**
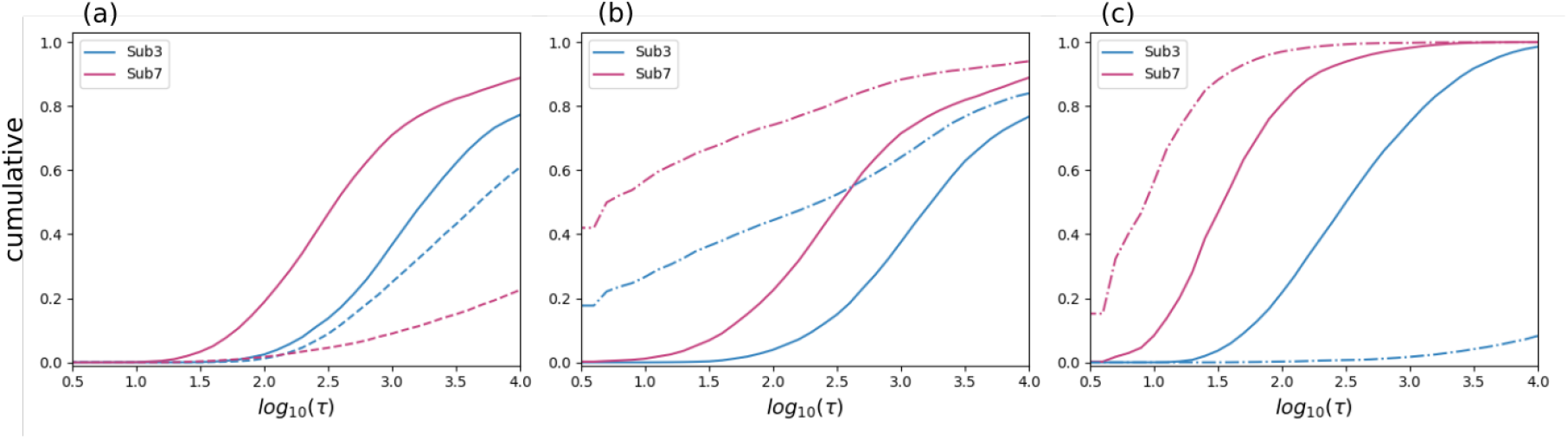
Speed inversion occurs also in the associative memory model. **(a)**Cumulative distribution for *τ* (on a log scale) without external input. Dashed curves are for the homogeneous network. **(b)**The input-driven divergence times, i.e., when half of the *S* = 3 active units are perturbed (*η* = 0.5, solid curves) and all of the *S* = 3 active units are perturbed (*η* = 1.0, broken curves). **(c)**Asymmetric connections between the two sub-networks, obtained by removing/pruning 30% of them, results in only quantitative changes. The slow-down and also the speed-up are dramatic, instead, when in addition, like in Fig.8c, the newly activated units are clamped by persistent external inputs (broken curves). For all panels, *T* = 0.05.

What happens when applying, after thermalization, an external input to some of the *S* = 3 units? Not much, Fig.9b shows, if the simulated input is applied to half of them (following the procedure used for Fig.8c, with *η* = 0.5 and no clamp). If *η* = 1.0, instead, i.e. the input is applied to the entire *S* = 3 sub-network, the there is a major effect, particularly in producing immediate or very early divergence of many of the trajectories, but the speed inversion remains more or less unaltered (broken lines).

Finally, Fig.9c shows that introducing moderate levels of asymmetry by diluting or cutting 30% of the connections *between* the two sub-networks does not have much of an effect either – unless one also clamps some of the units in the posterior network, in the manner already described; then, the posterior network slows down, almost to a standstill, which is intuitive, while surprisingly the anterior network speeds up further, as if unable to find any single satisfactory accomodation to the configuration imposed posteriorly.

## 4 Discussion

Our study is premised on the hypothesis that some of the characteristics of cortical dynamics have their roots in the statistical physics of disordered systems. Prior to attempting to validate the connection between two levels of analysis so distant from each other, we wanted to explore what the statistical properties might be, that might find – or not – their expression at the neural systems level. We have considered the reduction of Braitenberg’s model of cortical connectivity to a Potts network, and reviewed the thermodynamic analysis that predicts different types of transition from paramagnetic to glassy phase, as a function of the number *S* of local states. Surprisingly, when combining in a “hybrid” network two halves with “low” and “high” *S* units, the former are slowed down by the interaction, and the latter are sped up, to the point of overtaking the former. This effect might be related to the different order of the phase transition to the glassy phase, but remarkably it is a reversal of the difference presented by homogeneous networks. Although one can construct seemingly intuitive explanations *a posteriori*, those did not enable us to *predict* it, in the least.

The speed inversion effect appears to survive largely unaltered the introduction of additional elements and details, and, importantly, the replacement of the random network with an associative memory with connections structured by learning.

What are the implications for cortical processing? First, one should note that such implications should be taken with more than a grain of salt, if anything because the key concept of a single cortical axis is rather ill-defined, at best. Perusing the many parameters of cortical circuitry that have been observed to vary across cortical areas, and the many more likely to be reported in the future, describing their variation as aligned to an axis, let alone whether it is the *same* axis across parameters, is a wishful simplification. The sensory-motor hierarchies conceptualized e.g. by Fuster [40] have their final station in motor cortex *after* passing through the more anterior prefrontal cortices, while the termination layers of intracortical fibers, used to distinguish between feed-forward and feedback projections, define a cortical hierarchy with the hippocampus at the top, the limbic cortices next to it just below, then the association cortices of both temporal and frontal lobes, going down all the way to primary sensory *and* motor cortices [41]. In terms of the number of largely local inputs to the basal dendrites of pyramidal cells, instead, Elston [42] gives estimates for areas V1, 7a, TE and 12, in macaque monkey, that are roughly in the ratios 1:4:11:16, more or less along a posterior-to-anterior axis – but then measures in other areas do not necessarily align, for example area 10 at the frontal pole is anterior to 12, but its pyramidal cells are estimated to have on average 17% fewer spines.

Our hybrid Potts network discards such complexity anyway to favor simplicity, and the speeding up of the large-*S* units that it reveals may have to be factored in, as an underlying phenomenon, in any complex scenario that envisages an imbalance between the effective numerosity of local attractor states across the cortical mantle.

Interestingly, machine learning has pointed out the usefulness of combining “processing units” with memory properties at different time scales (*LSTM* units), e.g. to tackle syntax in language production and understanding. In particular, it has been predicted that long- and short-range units, which are taken to correspond to patches of cortex of perhaps 10^6^ neurons, similar to our Potts units, should reside in different cortical regions [43]. Our findings should prove useful to research in this natural language processing framework, by at least contributing a warning that the properties of the units in a homogeneous network, or even in isolation, may differ, to the point of being the opposite, from those of the same units in a hybrid one.

A rather different linguistic domain in which the effective speed or slowness of glassy dynamics may be important is language evolution. There, it has long been hypothesized that the syntactic parameters that determine the internal structure of language and that evolve or even “mutate”, like units of a genetic code, on a scale of hundreds or thousands of years [44], may all be *binary*. Notably, many other features which are needed to describe natural languages and to implement them in artificial systems are obviously far from binary and appear to evolve, largely, on faster time scales. Our study suggests that in a network of parameters with effectively random interactions, those that emerge in evolution as more resistant to change, and therefore describe the most stable internal structure or set of *motifs* of a natural language are precisely the binary ones, whether or not they possess a default value [45].

Yet other seemingly distant domains are those of protein folding and evolution, which have been approached with simplified Potts models [46, 47]. The possible application of our results to these different fields is left for future work.

## 5 Appendix

### 5.1 The first-step RSB free energy of Potts glass model

The Edward-Anderson order parameter is set as

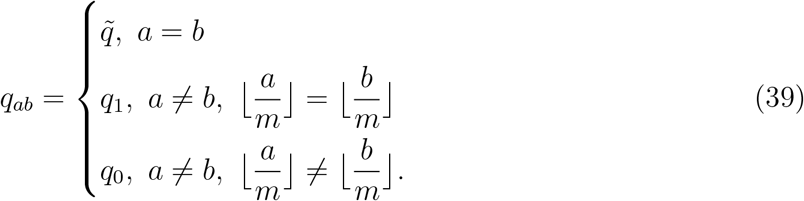

Then the free energy reads,

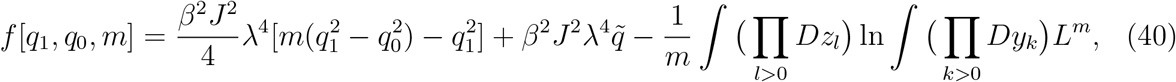

where

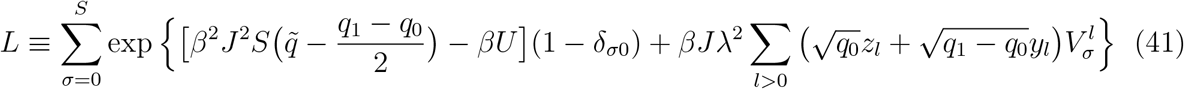

and

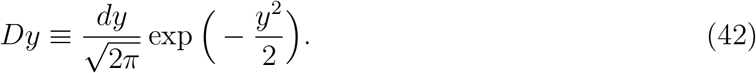

### 5.2 Finite number of patterns in associative memory

We compute the maximum value of *U*, below which the onset of glassy phase is continuous in the limit of *α* = *p*/*N* → 0. The trivial solution of Eqs. (28), 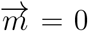, is stable as long as the corresponding eigenvalue

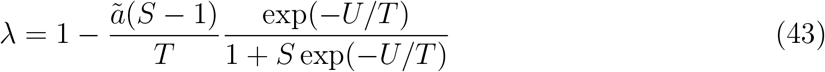

remains positive. This is always the case for *T* > *T_c_*, the critical temperature where the trivial solution becomes unstable and Mattis states appear. Mattis states satisfy

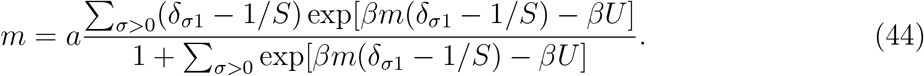

and the critical temperature for a given value of *U* is determined by solving

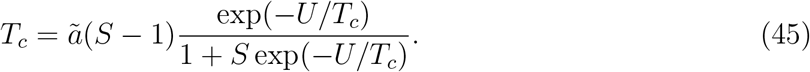

We can see that if *U* → +∞, *T_c_* → 0 and the trivial solution 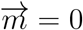 is stable for all temperature.

In Fig.10, we show values of *U_c_* and the critical temperature at *U* = *U_c_*.

**Figure 10:**
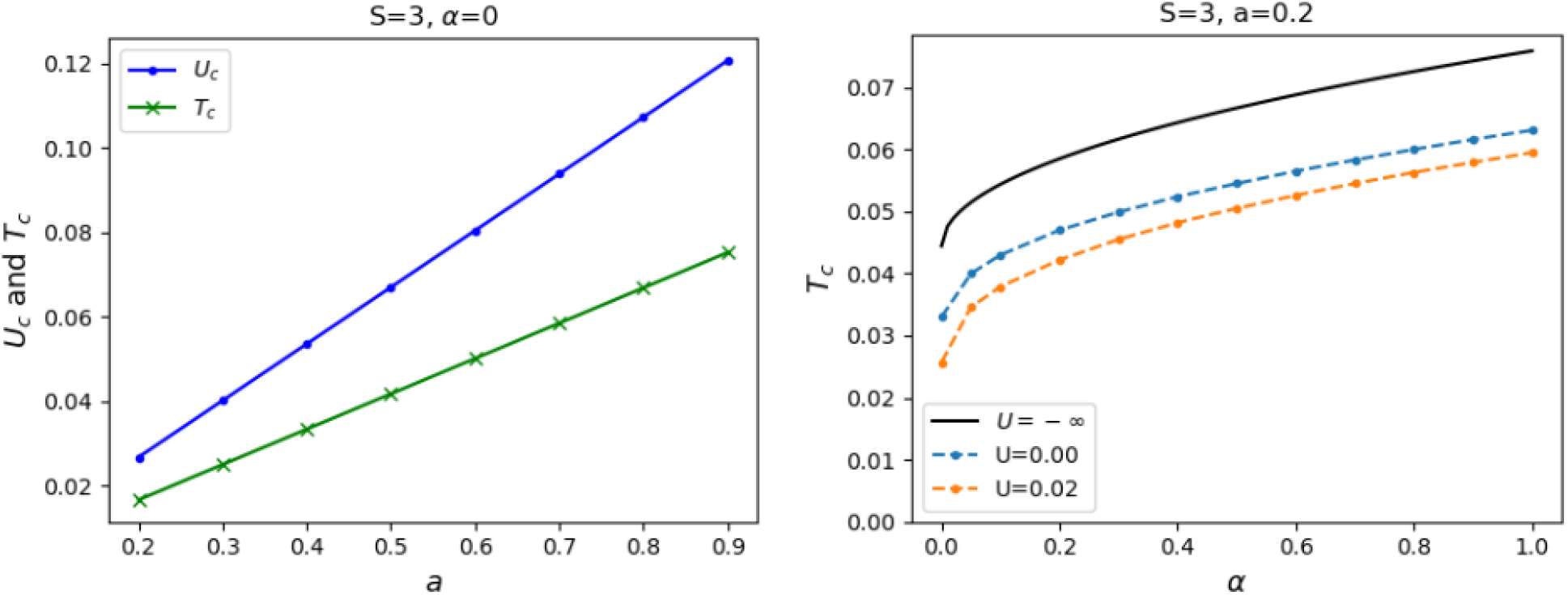
High temperature phase of associative memory for *S* =3. **Left**: maximum value of *U* (blue) above which the transition is no longer continuous, and the corresponding critical temperature (green) are plotted against sparsity of patterns for *α* → 0. **Right**: Critical temperature as a function of *α* for *α* = 0.2. Note that *U* = 0.02 is just below *U_c_* ≈ 0.026 given on the left panel.

### 5.3 Details on the computer simulations

For models with a quiet state, the Edward-Anderson order parameter is computed as, instead of Eq. (37),

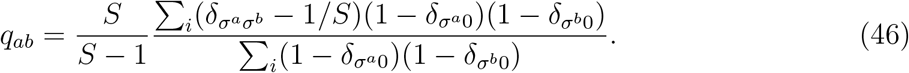

The mean activity of the network is controlled by time-dependent threshold

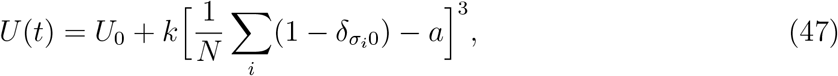

where *a* is the sparsity of patterns in associative memory model and *k* is set as 1000. For the Potts glass model with a quiet state, we have used the same activity level *a*.

The external input to the posterior sub-network is modelled by persistent external fields applied (after thermalization) to a fraction *η* of its units, which will maintain its states during dynamics (clamping in the main text). Specifically, we randomly select a fraction *η* of all active units in the *S* = 3 sub-network. Among the selected units, a fraction a of them is flipped into a different active state, while the remaining fraction 1 – *a* of them is set into a quiet state. The same number of units among quiet units is activated to maintain the same level of activity.

If not specified explicitly, parameters are set as in Table 1.

**Table 1:** Parameters of the network

| Symbol | Meaning | Default value |
| --- | --- | --- |
| $N$ | number of Potts units | 256 |
| $S$ | number of states per unit | 7 (3) |
| $T$ | temperature (noise level) | 0.5 |
| $\gamma$ | degree of asymmetry | 0.2 |
| $\eta$ | fraction of units with external inputs | 0.5 |
| $p$ | number of memory patterns | 1024 |
| $t_0$ | number of thermalization updates | 1000 |
| $a$ | mean activity | 0.25 |

## Notes

### Competing Interest Statement

The authors have declared no competing interest.

### Summary of Updates

Two panels of Fig. 7 (b and c) and two panels of Fig. 8 (a and b) are revised. A sentence in line 282-284 is revised to reflect the above changes in figures. Accordingly, one of default values in Table 1 is corrected. Except for these changes, everything remains the same.

